# Revisiting the Neural Architecture of Adolescent Decision-Making: Univariate and Multivariate Evidence for System-Based Models

**DOI:** 10.1101/2020.11.26.400416

**Authors:** João F. Guassi Moreira, Adriana S. Méndez Leal, Yael H. Waizman, Natalie Saragosa-Harris, Emilia Ninova, Jennifer A. Silvers

## Abstract

Understanding adolescent decision-making is significant for informing basic models of neurodevelopment as well as for the domains of public health and criminal justice. System-based theories posit that adolescent decision-making is guided by activity amongst reward and control processes. While successful at explaining behavior, system-based theories have received inconsistent support at the neural level, perhaps because of methodological limitations. Here, we used two complementary approaches to overcome said limitations and rigorously evaluate system-based models. Using decision-level modeling of fMRI data from a risk-taking task in a sample of 2000+ decisions across 51 human adolescents (25 females, mean age = 15.00 years), we find support for system-based theories of decision-making. Neural activity in lateral prefrontal cortex and a multivariate pattern of cognitive control both predicted a reduced likelihood of risk-taking, whereas increased activity in the nucleus accumbens predicted a greater likelihood of risk-taking. Interactions between decision-level brain activity and age were not observed. These results garner support for system-based accounts of adolescent decision-making behavior.

**Significance Statement:** Adolescent decision-making behavior is of great import for basic science, and carries equally consequential implications for public health and criminal justice. While dominant psychological theories seeking to explain adolescent decision-making have found empirical support, their neuroscientific implementations have received inconsistent support. This may be partly due to statistical approaches employed by prior neuroimaging studies of system-based theories. We used brain modeling—an approach that predicts behavior from brain activity—of univariate and multivariate neural activity metrics to better understand how neural components of psychological systems guide decision behavior in adolescents. We found broad support for system-based theories such that neural systems involved in cognitive control predicted a reduced likelihood to make risky decisions, whereas value-based systems predicted greater risk-taking propensity.

## Introduction

Adolescent decision-making has important implications for basic science (Blakemore & Mills, 2014; Larsen & Luna, 2018; Sharp & Wall, 2018; Yeager, Dahl, & Dweck, 2018) as well as public health, civic matters, and criminal justice policy (Cohen, Bonnie, Taylor-Thompson, & Casey, 2015; Cohen & Casey, 2014; Oosterhoff & Wray-Lake, 2020). Influential theories posit that adolescent decision-making is governed by psychological “systems” that compete (or in some cases, complement) to guide behavior (Casey, 2015; Shulman et al., 2016). While system-based theories have enjoyed broad success at describing the *psychological* underpinnings of adolescent decision behavior, they have yielded mixed findings when used to describe the *neurobiology* underlying said behavior (e.g., Pfeifer & Allen, 2012). This discrepancy between psychological and neural data may be due in part to prior neuroimaging work employing brain mapping (predicting brain from behavior) instead of brain modeling (predicting behavior from brain) approaches, testing theory between- instead of within-subjects, and not considering multivariate neural patterns. The current neuroimaging study sought to overcome these methodological limitations, and to more rigorously test the validity of system-based models for predicting adolescent risky decision-making.

A number of system-based theories have been used to explain risky decision-making and related motivated behaviors in adolescence (Casey, 2015; Ernst, Pine, & Hardin, 2006; Strang, Chein, & Steinberg, 2013). Most of these theories contain two key elements. First, they posit the existence of two (though some posit three) adversarial systems: A value-based system oriented toward immediate incentives, increasing the propensity for risk-taking, and a cognitive control system that restrains the former system to avoid risks. Second, these prominent theories argue the value-based system is primed to ‘overpower’ the cognitive-control system in adolescence (i.e., they interact with age), ostensibly leading adolescents to take more risks than children and adults – particularly in socioemotional contexts (Shulman et al., 2016; Steinberg et al., 2017). System-based models tend to perform well at explaining adolescent behavior in observational and experimental studies (Botdorf, Rosenbaum, Patrianakos, Steinberg, & Chein, 2016; Ellingson, Corley, Hewitt, & Friedman, 2019; Steinberg et al., 2017). However, *neuroscientific evidence* for these theories is far less consistent (Flannery et al., 2017; Lee et al., 2018; van Duijvenvoorde, Achterberg, et al., 2016; van Duijvenvoorde, Peters, et al., 2016), prompting calls to update system-based theories (Casey, 2015; Pfeifer & Allen, 2012, 2016), or revise them so drastically as to be categorically different from existing versions (Harden et al., 2017; Romer et al., 2017). Without outright rejecting these possibilities, we propose an alternative interpretation for why system-based theories receive inconsistent neuroscientific support.

Most prior neuroscientific investigations of adolescent decision-making have relied on univariate *brain mapping* methods to compare individuals who differ in terms of age or risk-taking behavior. Brain mapping refers to statistically predicting brain activity from stimulus or task characteristics, or task behavior (Kragel, Koban, Barrett, & Wager, 2018). An alternative to brain mapping is *brain modeling* (Kragel et al., 2018), which uses neuroscientific data to predict cognitions and behavior (i.e., any kind of neural metric predicting behavior). Having recently grown in popularity, brain modeling approaches have seen broad applications, some of which involve within-person modeling, including prediction of food craving (Cosme & Lopez, 2020; Cosme, Ludwig, & Berkman, 2019), emotion regulation tendencies (Doré, Weber, & Ochsner, 2017), negative affect (Chang, Gianaros, Manuck, & Krishnan, 2015), chronic pain (Wager et al., 2013) and vision (Gardner & Liu, 2019; Liu, Cable, & Gardner, 2018). While brain mapping has generated key discoveries in neuroscience (e.g., Kanwisher, 2017), it can be problematic for evaluating system-based theories. Statistically, brain modeling may be preferable to brain mapping because individual units of analysis (e.g., voxels or neurons) are more predictive when used in concert (such as in a multivariate signature) to predict task behavior, as opposed to the opposite (e.g., behavioral responses predicting brain activity) (Zhao et al., 2020). That is, treating individual voxels as the outcome of an analysis is less informative than examining how multiple voxels collectively predict a phenomenon of interest. Unfortunately, prior brain-mapping studies testing system-based theories of decision-making have largely overlooked the cumulative information that comes from many voxels. Another advantage of brain modeling is that it is better suited for trial-level, within-subject modeling, which tends to be better powered than classic between-subject analyses. Philosophically, system-based theories make predictions about how underlying neural processes drive behavior – for example, “when value-based brain activity is high, individuals will be more likely to take a risk” – which almost by definition aligns with brain modeling. Relatedly, system-based theories of decision-making are implicitly geared towards explaining *within*-subject behavior (Strang et al., 2013), yet most prior studies of adolescent risky decision-making have focused on *between*-subject differences (Flannery et al., 2017; Rudolph et al., 2017; van Duijvenvoorde, Achterberg, et al., 2016). Understanding within-adolescent fluctuations in decision-making carries critical translational implications for understanding why the same individual may be law-abiding most of the time but occasionally engage in destructive or maladaptive behavior. The aforementioned limitations of prior work motivated the present study to employ novel methodology to test the validity of system-based accounts for predicting adolescent decision making.

## Methods

### Overview

The current study is, to the best of our knowledge, the first within-subject, brain modeling investigation of system-based theories of adolescent decision-making. Using functional magnetic resonance imaging (fMRI), we predicted trial-by-trial risky decision-making in healthy adolescents as a function of brain activity from value-based and cognitive control systems, the first premise posited by system-based models. We then tested to see if the two systems interacted with age, testing the second premise posted by system-based models. Critically, we examined two versions of value-based and cognitive control systems: a ‘classic’ univariate version and a newly posited ‘switchboard’ multivariate version. The classic variant of the theory assumes that the value-based system that prioritizes immediate rewards is primarily housed in the nucleus accumbens (NAcc) whereas the cognitive control system is located in lateral prefrontal cortex (lPFC) (Shulman et al., 2016). This variant is clearly modular, in that it posits that psychological functions are represented in isolated brain regions, or modules, that independently and locally perform their respective function. While evidence exists to suggest that some degree of modularity may be present in the brain (Kanwisher, 2017), this assumption is inconsistent with much other work in cognitive neuroscience that shows psychological processes are encoded in distributed, multivariate signatures (Chang et al., 2015; Huth, Heer, Griffiths, Theunissen, & Gallant, 2016; Parkinson, Kleinbaum, & Wheatley, 2017). To that end, we additionally tested a ‘switchboard’ version of the model wherein we predict risky behavior as a function of multivariate neural signatures of value and cognitive control (via the use of multivoxel pattern analysis; MVPA). The advantage of this approach is that it does not hypothesize the localization of mental function to any given region of interest (ROI) but instead assumes that mental functions are encoded in distributed patterns. Another way to summarize the two variants of the model is that the functional units of the classic model lie in particular ROIs, whereas the functional units of the switchboard model are comprised by patterns of activity that cut across brain regions. We used multilevel logistic regression to examine how linear combinations of these brain metrics predicted *within-person* risky behavior. Last, for thoroughness we also implemented a between-subjects version of the brain model (predicting risky behavior as a function of brain metrics using only between-subjects information) while considering between-subject variables including age and sex as predictors, in addition to a traditional univariate analysis.

### Procedures and Measures

#### Participants

The *N* = 51 participants (Mean age = 15.00 years, SD = 3.66, range = 9.11 - 22.60, 25 females) in the current study were part of a broader longitudinal study investigating the impact of early life experiences on the neural bases of socioemotional development. This age range is consistent with recent scientific advances that suggest adolescence in human development may last nearly fifteen years (Kinghorn, Shanaube, Toskas, Cluver, & Bekker, 2018). Participants in the current set of analyses were those who provided usable data from an fMRI scanning session and did not have a history of early social deprivation. Ethnically, eight participants identified as Hispanic/Latinx (15.7%). Racially, twenty-six participants identified as white (51%), six participants (11.8%) identified as Asian/Asian American, one participant (2%) identified as Native Hawaiian/Other Pacific Islander, seven participants (13.7%) identified as African American, no participants (0%) identified as Native American/Alaskan Native, four participants (7.8%) identified as being mixed race, four participants identified as belonging to an unlisted race (7.8%), and three participants (5.9%) declined to report their race. Sample size was dictated by the number of participants willing to participate in this wave of data collection. Participants were compensated $50 (USD) for participating in fMRI scanning. The research was completed at the University of California, Los Angeles (UCLA). All participants under 18 years provided informed assent and their parents provided informed consent; all participants 18+ years provided informed consent. All research practices were approved by the Institutional Review Board at the University of California, Los Angeles. Data and analysis code are publicly available on the Open Science Framework (OSF; osfi.io/fphn4).

### Experimental Design

#### Risky Decision-Making Paradigm

Participants completed the Yellow Light Game (YLG) while undergoing fMRI scanning (Figure 1A; Op de Macks et al., 2018). An adaptation of a widely used adolescent risk-taking task (the stoplight game; Chein, Albert, O’Brien, Uckert, & Steinberg, 2011; M. Gardner & Steinberg, 2005; Peake, Dishion, Stormshak, Moore, & Pfeifer, 2013), the YLG is a computerized driving simulation in which participants drive along a straight road and encounter a series of intersections. Consistent with prior studies, participants in our study were told the objective of the game was to drive through the set of intersections as quickly as possible. The traffic light at each intersection turned yellow for 1000ms prior to crossing each intersection and participants were faced with a choice to brake (‘stop’) or drive through the intersection (‘go’). A choice to brake at the intersection resulted in a delay of 2500ms. A choice to accelerate through the intersection resulted in one of two outcomes -- (i) participants would drive straight through the intersection with no delay, or (ii) a car from the cross-street would crash into them resulting in a 5000ms delay. A 10000ms delay was imposed if participants failed to respond on a trial. Participants made their choices by pressing one of two buttons on a button box using their index and middle fingers.

**Figure 1.**
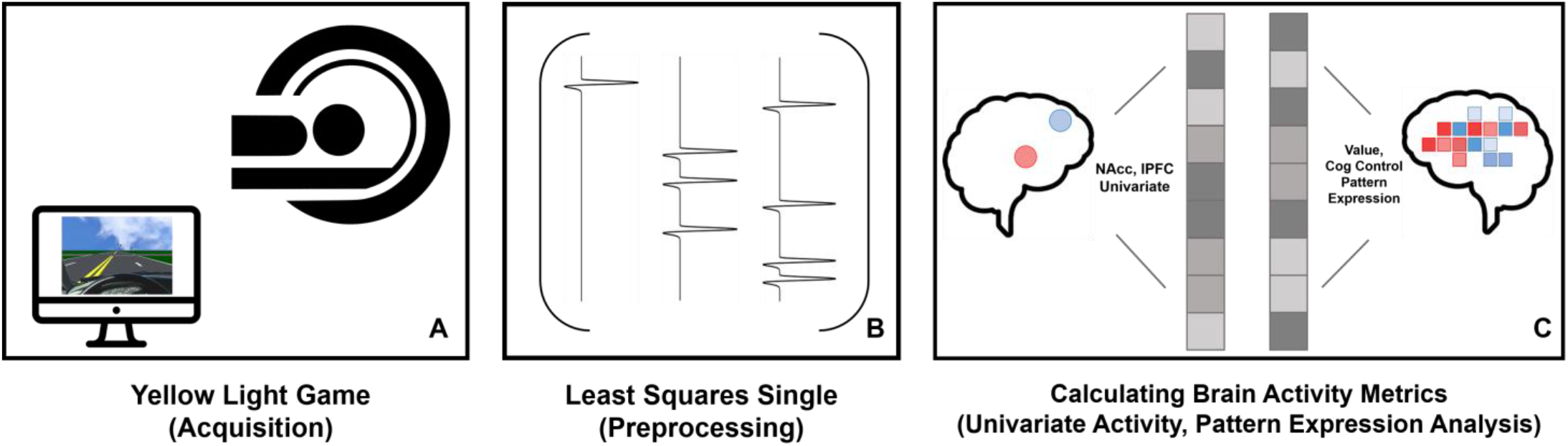
Schematic of Data Collection, Processing, and Analysis. Figure 1-1 visualizes head motion metrics for fMRI data. *Note.* Panel A depicts data acquisition of the Yellow Light Game while participants were undergoing fMRI scanning. Panel B depicts the Least Squares Single (LSS) modeling implemented as a preprocessing step. Panel C is a schematic of extracting the set of univariate and multivariate metrics from Panel B’s resulting beta-series.

Participants completed three runs of the task, consisting of 15 trials each (*n* = 45 total trials). Unbeknownst to participants, five intersections per run were set to result in a crash if participants chose to accelerate through them, meaning that the probability of crashing was equal to ⅓. Participants were not made aware of this probability. Each run had specific intersections that were rigged to crash and the order in which runs were administered was counterbalanced across participants. Buttons indicating ‘go’ and ‘stop’ were also counterbalanced between subjects amongst the index and middle fingers. The task was self-paced but typically took participants approximately 2.5 minutes to complete each run. Participants completed two, 10-trial practice runs prior to scanning in order to eliminate any potential confounds associated with learning. The YLG was programmed in Java and ran off Apache Tomcat, a program that creates a HTTP web-server environment.

#### fMRI Data Acquisition

Imaging data were acquired on a 3T Siemens Prisma scanner using a 32-channel head coil and a parallel image acquisition system (GRAPPA). A high resolution T1-weighted, magnetization-prepared rapid acquisition gradient echo (MPRAGE) image was acquired for registration to functional runs (TR = 2400ms, TE = 2.22ms, flip angle = 8°, FoV = 256mm^2^, 0.8mm^3^ isotropic voxels, 208 slices). Functional images were acquired using a T2^*^ EPI BOLD sequence. Thirty-three axial slices were collected with a TR of 2000ms and a 3 × 3 × 4 mm^3^ voxel resolution (TE = 30ms, flip angle = 75°, FoV = 192mm^2^). Participants completed the YLG by using a head-mounted on the coil to view an LCD back projector screen.

### fMRI Analysis

The following sections describe our approach to analyzing the fMRI data using both univariate activity estimates of the NAcc and lPFC and multivariate pattern expression values for reward and cognitive control signatures (signature definitions described below). We first describe our preprocessing steps and then outline the single trial analysis procedure used to produce both univariate activity estimates and multivariate metrics for each trial during the task. Because we were interested in within-subject variability in decision-making, we estimated univariate and multivariate values for each trial across all subjects. Single-trial metrics were used for both the within-person (in disaggregate form) and between-person (in aggregate form) analyses, for consistency. We also detail how we conducted the traditional univariate analysis of the YLG.

### Preprocessing

Prior to preprocessing, functional images were visually inspected for artifacts and biological abnormalities. No images contained obvious artifacts or biological abnormalities that warranted exclusion from further analysis. fMRI data were preprocessed and analyzed using the fMRI Expert Analysis Tool (FEAT, version 6.00) of the FMRIB Software Library package (FSL, version 5.0.9; fsl.fmrib.ox.ac.uk). Preprocessing consisted of the following steps: We used the brain extraction tool (BET) to remove non-brain tissue from functional and structural runs, spatially realigned functional volumes to the middle image to correct for head motion using MCFLIRT, and high-pass filtered the data with a 100-s cutoff. We used fsl_motion_outliers to identify volumes that exceeded a 0.9mm frame displacement (FD) threshold for head motion (Siegel et al., 2014), though most participants failed to record any volumes exceeding this threshold (Table 1-1; Figure 1-1). No participant had more than 10% of their volumes in a given run exceed the aforementioned framewise displacement threshold and thus, no participants were excluded on the basis of head motion in our sample. Spatial smoothing was not conducted during preprocessing and was instead applied later when extracting data from single trial activity estimates because the extent of smoothing depended on the type of information that was being extracted from the single trial (average ROI activation warrants greater smoothing than pattern expression analysis). We prewhitened the data to correct for autocorrelated residuals across time. Functional data were registered to each subject’s high resolution MPRAGE scan with FSL’s boundary-based registration (Greve & Fischl, 2009) while maintaining the 3 × 3 × 4 mm voxel size. To preserve the fine-grained spatial resolution of the data, we did not register the functional runs to standard MNI space at this stage but did so for the traditional group analyses (See Table 7). As detailed below, masks and neural signatures were defined in standard space and then transformed to subject space.

### Within-Subject Analyses

#### Single Trial Activity Estimation

We used a least squares analytic framework to obtain trial-level estimates of the BOLD signal (i.e., a beta-series; Rissman, Gazzaley, & D’Esposito, 2004). Here we opted to use the least squares single (LSS; Figure 1B) method, due to its advantageous statistical properties over the least squares all (LSA) estimator, especially considering the fast timing of the YLG (Mumford, Davis, & Poldrack, 2014; Mumford, Turner, Ashby, & Poldrack, 2012). Accordingly, a fixed-effects General Linear Model (GLM) was created for the decision period on each trial of the YLG game within each participant. A decision period was defined as the time between the onset of the yellow light (i.e., when the light at the traffic intersection switched from green to yellow as the car approached the intersection) and when participants pressed a button to signify their decision. A GLM was modeled for the *i*-th decision period (target decision) such that the target decision received its own regressor, all other decision periods were modeled in a single, separate nuisance regressor, and outcomes of all decisions (delays due to the braking, successful passes after running the light, or crashes) were modeled in another regressor.

Head motion was statistically controlled for across all GLMs by adding FSL’s extended motion parameters (6 regressors for x, y, z, pitch, roll, yaw directions, their squares, and their derivatives, comprising 24 regressors) in addition to regressors for single volumes that exceeded a frame displacement threshold of 0.9mm (i.e., censoring). The first temporal derivative of all task and motion regressors were also entered into the model in order to account for slice timing and motion effects, respectively. Parameter estimates from each trial-specific GLM were used to create a linear contrast image comparing the target decision period to the implicit baseline (unmodeled events). We then used the unthresholded z-statistics of this contrast to extract univariate and multivariate estimates of the BOLD signal in regions of interest.

#### Extracting Univariate ROI Activity from Single Trial Estimates (Classic Model)

Masks were defined to extract univariate activity from the NAcc and lPFC. Both masks were defined using the Harvard-Oxford probabilistic atlas as rendered in FSL’s viewer (fslview) on the MNI152 NLIN 6th generation T1 template image at 2 mm^3^ voxel resolution (avg152T1_brain.nii.gz; Brett, Johnsrude, & Owen, 2002). This atlas contains probabilistic masks to various bilateral structures that articulate the probability that a given voxel within the mask falls in the specified brain region. We created a bilateral NAcc mask by merging the atlas’ left and right nucleus accumbens probabilistic images into a nifti volume and thresholding the image at *p* = .25. We selected the nucleus accumbens due to prior empirical and theoretical accounts of this region’s importance in adolescent risk-taking (Galvan et al., 2006; Steinberg, 2010). The .25 threshold was selected with the goal of creating a mask that was relatively inclusive but did not also possess clear outlying voxels (i.e., voxels with a very low probability of landing in the accumbens). A similar procedure was used to create a bivariate lPFC mask by selecting and merging left and right interior frontal gyrus masks (both the pars opercularis and pars triangularis) and thresholding the image at *p* = .50. We chose a .50 threshold for this mask because lPFC activation reported in prior adolescent neuroimaging studies tends to be spatially broad. However, we also created another version of this mask by thresholding at *p* = .25 in order to be consistent with the NAcc mask and found broadly consistent results (masks shown in Figure 2).

**Figure 2.**
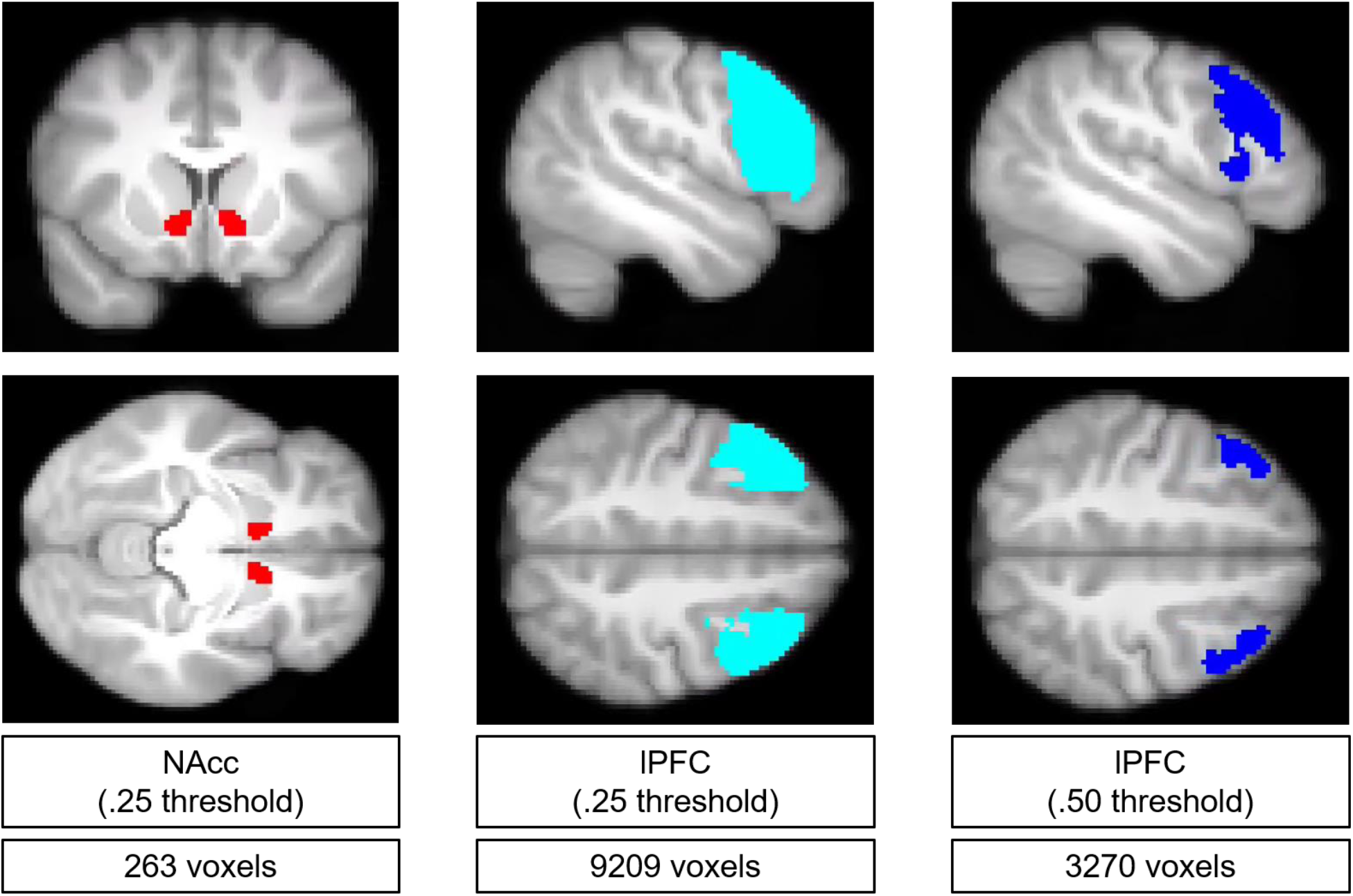
Nucleus Accumbens and Lateral Prefrontal Cortex Masks Used to Extract Univariate Activation Estimates *Note.* NAcc refers to ventral striatum (NAcc); lPFC refers to lateral prefrontal cortex. Thresholds were applied to probabilities values from the Harvard-Oxford cortical and subcortical atlases. Masks are depicted in MNI standard space and projected onto an average of all subjects high resolution anatomical images.

Once our masks were defined, we transformed the masks into the native space for each single trial activity map using FLIRT (i.e., whole brain zstat) and then extracted activity estimates using the nilearn software package (Abraham et al., 2014). We used the package’s `NiftiMasker()` function to mask each single trial activity estimate with the aforementioned NAcc mask and then again with the aforementioned lPFC mask and extract the mean of all voxels within each respective mask (Figure 1C). It was at this step that we applied smoothing to the extracted data (6 mm, fwhm), as the `NiftiMasker()` function allows one to smooth a masked image when extracting data. This step produced a set of NAcc and lPFC activation estimates for each trial on the task across all subjects (i.e., each subject had as many NAcc and lPFC activation estimates as they did decisions in the YLG).

#### Computing Pattern Expression from Single Trial Estimates (Switchboard)

We used pattern expression analyses to quantify the extent to which whole-brain patterns of brain activity corresponded to neural signatures of cognitive control and value-based computations (Figure 1C). Such an analysis allows one to determine how strongly a given pattern of brain activity is expressed as a function of a neural signature of interest (Chang et al., 2015; Kragel et al., 2018; Wager et al., 2013). Neural signatures are thought to be the fingerprints of brain activity that encode a particular psychological process or state of interest. In practice, they are frequently defined as maps of the brain containing weights that quantify the strength and direction of association between each voxel and the psychological process of interest.

The first step in this analysis involved defining neural signatures of cognitive control and value-based computation. To this end, we used Neurosynth, a web-based platform that automates meta-analysis over a large set of published fMRI studies (Yarkoni, Poldrack, Nichols, Van Essen, & Wager, 2011), to retrieve meta-analytic maps (uniformity) for the terms ‘value’ (470 studies) and ‘cognitive control’ (598 studies). We chose these terms based on system-based theories such that ‘value’ references one system which drives adolescents to make risky decisions in service of acquiring immediate hedonic rewards whereas ‘cognitive control’ references a second system which modulates the drive towards immediate rewards (Shulman et al., 2016; Steinberg, 2013). A benefit of using meta-analytic maps as neural signatures is that they ‘allow the data to speak for themselves’ by allowing us to select voxels weights that are most strongly associated to our psychological processes of interest (in contrast to approaches that posit singular ROIs that might exclude meaningful voxels). To our knowledge, the majority of data used to calculate these analytic maps come from traditional univariate studies, though we note that the high volume of studies should theoretically allow for identification of the most sensitive voxels. To ensure the robustness of results, we used both the uniformity and association maps (reported in Table 3). While a review of the differences between these two types of images is beyond the scope of this paper (see Neurosynth.org/faq), we briefly note here that association maps provide greater selectivity about the relationship between a voxel and a given term by incorporating information about base rates. To be comprehensive, we re-ran all analyses with the Neurosynth term ‘reward’ and observed nearly identical results. Maps of the two signatures (uniformity) are depicted in Figure 3.

**Figure 3.**
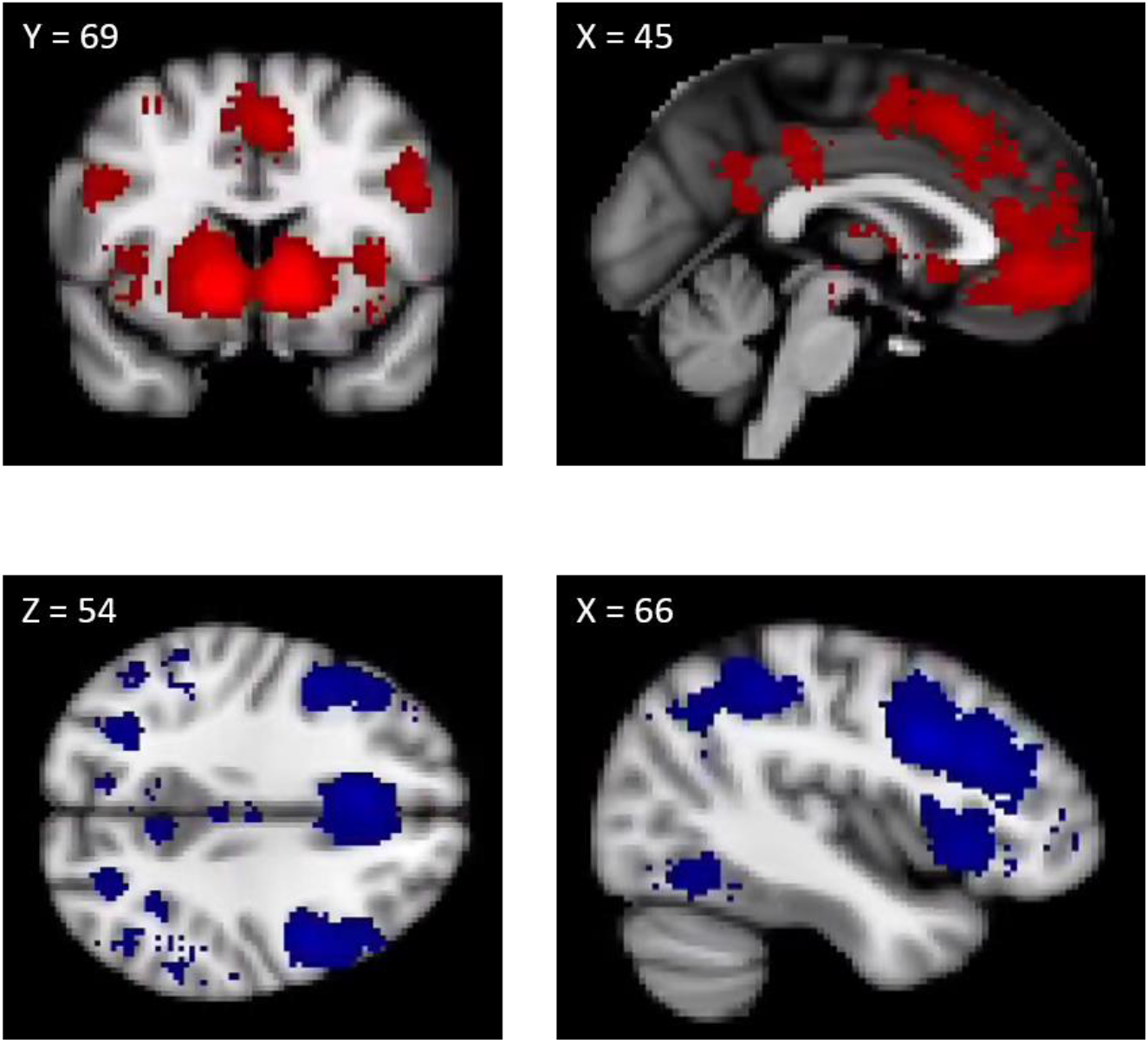
Multivariate signatures of Value (top row) and Cognitive Control (bottom row) *Note.* Both signatures obtained from Neurosynth. Uniformity signatures are depicted here. Voxels weights differed between each mask (i.e., a hypothetical voxel could be included in both signatures, but its weight likely varied between signatures. This is important to note because these maps were used as multivariate signatures, which ultimate meant that the same brain regions included in both masks possessed a different multivariate signature.

Once the neural signatures were defined, we transformed each signature into the native space of each single trial activity map (i.e., whole brain zstat), and extracted multivariate patterns from both the transformed neural signatures and activity estimates using `NiftiMasker()`. Multivariate patterns were minimally smoothed (1mm fwhm; Weaverdyck, Lieberman, & Parkinson, 2020) and then the dot product between voxels in the two patterns (activity estimate, neural signatures) was taken (we re-ran all analyses with a greater smoothing kernel–4mm fwhm–and obtained highly similar results). This resulted in two pattern expression estimates per trial, one quantifying the expression of value patterns in brain activity during a given decision and another quantifying the expression of cognitive control patterns in brain activity during the same decision. Barring missing decision data (see Figure 4), each subject had 90 pattern expression estimates − 45 for value and 45 for cognitive control, each corresponding to a decision during the yellow light game.

**Figure 4.**
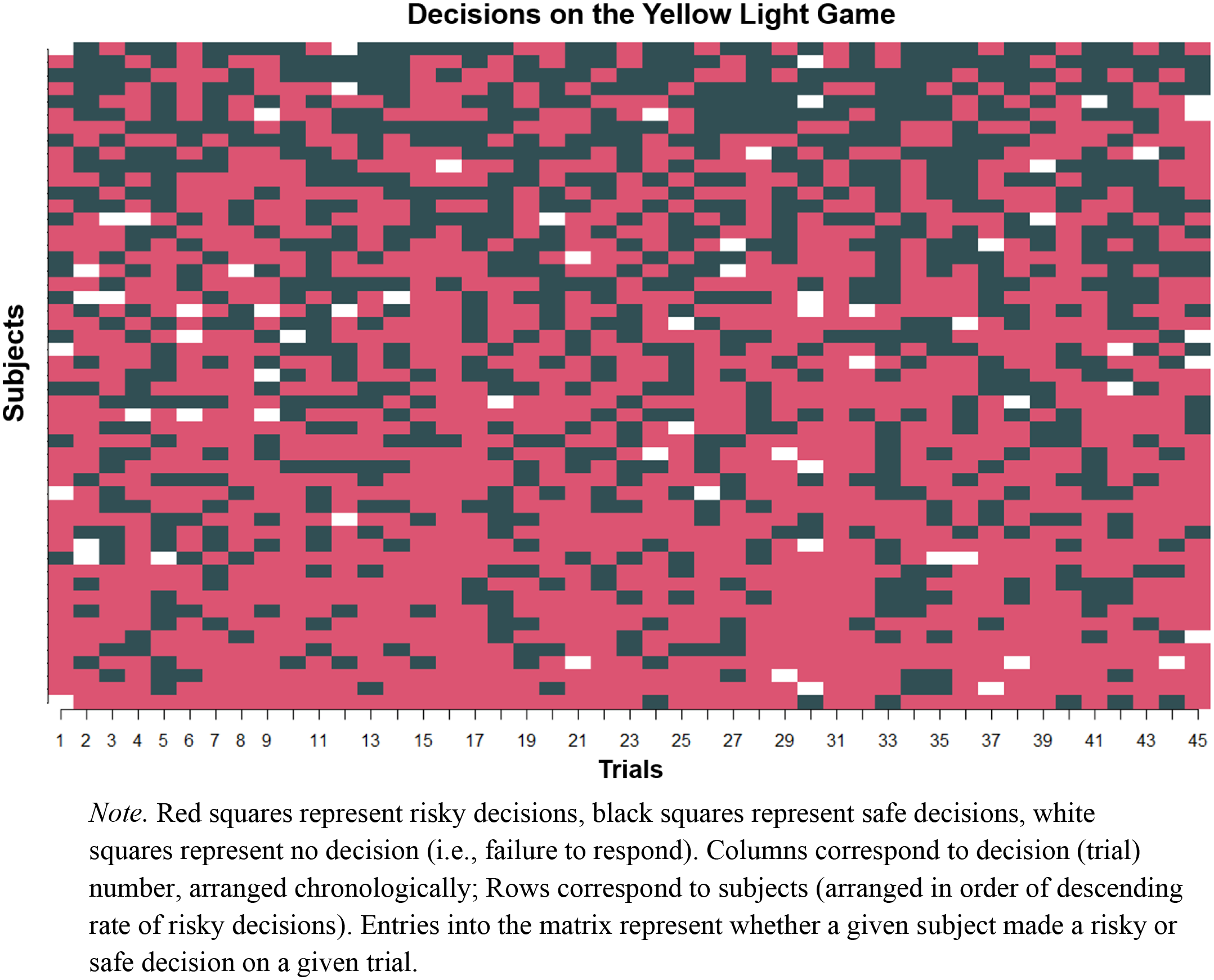
Visualizing risky and safe decisions on the yellow light game for all participants *Note.* Red squares represent risky decisions, black squares represent safe decisions, white squares represent no decision (i.e., failure to respond). Columns correspond to decision (trial) number, arranged chronologically; Rows correspond to subjects (arranged in order of descending rate of risky decisions). Entries into the matrix represent whether a given subject made a risky or safe decision on a given trial.

Notably, we were aware of previous work using pattern expression analyses with a preprocessing stream that involved normalizing images to standard space (Chang et al., 2015; Wager et al., 2013). We note that our decision to keep images in subject space for pattern expression calculation is not necessarily at odds or incompatible with these prior studies, best practices for pattern expression analyses, or even the broader MVPA literature. Unlike these prior studies, our goal was not to create a biomarker or construct a neural signature that can be applied across an entire population (Weaverdyck et al., 2020). Because our focus was on intra-individual fluctuations in activity and links to decision-making behavior, it was appropriate to refrain from normalizing to preserve fine-grain spatial information.

### Between-Subject Analyses

#### Aggregation of Trial-Level Data for Between-Subjects Analysis

In order to conduct between-subject analyses, we aggregated the trial-level univariate and pattern expression data. We did this by taking the average of the aforementioned brain activity metrics for each subject.

#### Group-Level Brain Mapping Analysis

We conducted traditional, group-level brain mapping (mass univariate) analyses to serve as a comparison point and complement our between-subject analyses (Chein et al., 2011; Op de Macks et al., 2018; Telzer, Ichien, & Qu, 2015). To this end, we first submitted each participant’s run-level data to a fixed GLM analysis in FSL. For this purpose, the YLG was modeled consistent with other prior univariate studies by including a regressor for ‘Go’ decisions, a regressor for ‘Stop’ decisions, and a regressor for outcomes (regardless of type, e.g., successful pass, crash). This differed from the LSS analysis in that all events from each condition of interest (‘Go’ decisions, ‘Stop’ decisions, outcomes) are put into a single regressor for that condition, whereas the LSS analysis assigns a target decision trial (regardless of type) its own regressor, and all other decisions and outcomes are modeled as two separate nuisance regressors. The same pre-processing decisions steps were taken as in all other analyses (e.g., slice timing correction via adding temporal derivatives, adding extended motion parameters, censored volumes, etc.). The only exception was that we smoothed our data for this model (6mm, fwhm), and non-linearly registered high resolution anatomical images to the MNI152 template image (10 mm warp resolution), and used the subsequent transformation matrix to register the functional images to standard space.

Parameter estimates from this GLM were used to create linear contrast images comparing the ‘Go’ and ‘Stop’ conditions (‘Go’ - ‘Stop’, ‘Stop-Go’). Random effects, group-level analyses were performed on this contrast using FSL’s FLAME1 module. A cluster defining threshold of Z = 3.1 was used in conjunction with a familywise error rate of *p* < 0.05 and Random Field Theory cluster correction to address the problem of multiple comparisons. An additional whole-brain analysis regressed age (mean centered) on these contrasts but found no age effects.

### Statistical Analysis

#### Overview

Our analytic approach consisted of two parts. The first set of analyses examined neural systems underlying within-subject variability in decision-making, using both classic and switchboard models. The second set of analyses examined between-subject variability in decision-making. Here, we again compared classic and switchboard dual-systems models. A detailed description of both approaches follows below. For thoroughness, we also report a traditional between-subjects univariate analysis of the YLG in Table 7.

#### Within-Subjects: Modeling Trial-Level Influences of Brain on Behavior

We executed our within-subjects test of the classic and switchboard models with a series multilevel logistic regression models. For each theory (classic, switchboard), we conducted a multilevel logistic regression model including trial-level estimates of brain activity and subject level controls (age, gender). The form and specification of the statistical models for both variants follow.

Trial-level, Classic:

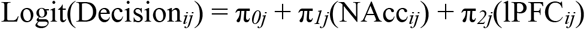

Trial-level, Switchboard:

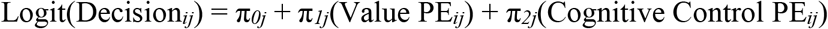

Decisions (1 = risky (‘Go’), 0 = safe (‘Stop’)) at the *i*-th trial for the *j*-th individual were modeled as a function of a subject-specific intercept (π_*0j*_), and brain activity metrics. The brain activity metrics in the classic model were average activations in the ventral striatum (NAcc_*ij*_) and lateral prefrontal cortex (lPFC_*ij*_) at the *i*-th trial for the *j*-th individual. Said activity metrics in the switchboard model were pattern expression (PE) estimates for value (Value PE_*ij*_) and cognitive control (Cognitive Control PE_*ij*_) for the *i*-th trial for the *j*-th individual. Subject-specific parameters for all within-person predictors (π_*1j*_ & π_*2j*_) correspond to the subject-specific expected change in the log odds of making a risky decision given a one unit increase in the predictor (e.g., average univariate brain activity, pattern expression score) holding the other predictors constant. All trial-level predictors were standardized using the grand mean. Re-running main analyses while standardizing trial-level predictors within-person produced statistically significant results with comparable parameter estimates (magnitude and sign).

As noted above, we controlled for the following between subject variables: gender (dummy coded, 0 = male, 1 = female) and age. The form of the between-subjects component of the model for both classic and switchboard follows (i.e., this component of the model was the same for both classic and switchboard models).

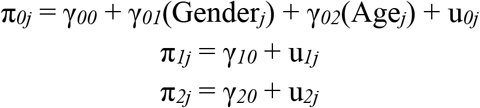

This component of the model reflects how all trial-level parameters are allowed to vary randomly between subjects (indicated by the u_*j*_’s, random effects) while showing the main effect of between subject predictors (γ_*01*_ & γ_*02*_). The other gammas in the model (γ_*10*_ & γ_*20*_) represent the fixed effect of the trial-level predictors (i.e., the portion of trial-level effects that are common to all participants). Random coefficient regression models were implemented with the ‘lme4’ package in R (Bates, Mächler, Bolker, & Walker, 2014) and significant tests were obtained using the ‘lmerTest’ package (Kuznetsova, Brockhoff, & Christensen, 2017). Here we note that this analytic framework affords us greater statistical power than we would focusing on a model exclusively testing between-subjects differences because we have many decisions nested within individuals. Because our predictors of interest occurred at the level of the decision, we were able to reach approximately 80% statistical power to detect a meaningful trial-level effect (Astivia, Gadermann, & Guhn, 2019; Schoeneberger, 2016). We also tested permutations of these models that allowed age to interact with the trial-level brain activity metrics, effectively testing the possibility that the strength of the two neural systems changes with age.

#### Modeling Between-Subject Brain-Behavior Associations

Using the aggregated univariate activity (i.e., the average of each subjects’ trial-level brain activity metrics) and aggregated pattern expression estimates (obtained via averaging over estimates within each subject), we sought to examine between subject brain-behavior associations. To do so, we conducted two multiple regression analyses. The first analysis examined the contribution of univariate NAcc and lPFC activity on the percentage of risky decisions during the task, while controlling for age and gender. The second analysis swapped out the univariate predictors for the multivariate pattern expression metrics.

## Results

### Baseline Models and Descriptive Data Visualizations

Individual decisions across all subjects are plotted in Figure 4. This figure highlights the variability in risky behavior both within and between subjects. Additionally, we ran two ‘baseline’ multilevel logistic regression models on the trial-level risky decision-making data from the YLG. The first model was an empty model, modeling trial-level decisions only as a function of an intercept, effectively estimating the unconditional likelihood of making a risky decision on the task (Table 1). The second model included gender and age as predictors (‘covariate-only model’) so as to examine the effects of these variables unconditioned on the brain activity data (Table 1)—neither were related to risky decision likelihoods.

**Table 1.** Log-odds of risky choice from Empty and Covariate-Only models. Table 1-1 lists head motion statistics for fMRI data.

| Term | Empty | Covariate-Only |
| --- | --- | --- |
|  | Estimate |  |
| Intercept | 0.527 (0.087)*** | 0.599 (0.119)*** |
| Trial Number | - | -0.009 (0.047) |
| Age | - | 0.104 (0.085) |
| Gender | - | -0.144 (0.169) |
| Variance Component | Estimate |  |
| Var(Intercept) | 0.282 | 0.268 |
| Var(Trial Number) | - | 0.006 |
| Fit Statistic | Statistic |  |
| AIC | 2887.1 | 2892.8 |
| BIC | 2898.5 | 2932.8 |

*Note.* ‘ refers to $p < .10$ , \* refers to $p < .05$ , \*\* refers to $p < .01$ , \*\*\* refers to $p < .001$ . Sex was dummy coded (0 = male, 1 = female). Standard errors of parameter estimates are printed in parentheses. Var() refers to a variance component of a given random effect from the model.

### Within Subjects Results

Results from our within-subject models are summarized in Tables 2-4, and Figure 5. Each are described in greater detail below.

**Table 2.** Log-odds of risky choice from within-subjects analysis of classic system-based models

| Term | A – Estimate | B – Estimate |
| --- | --- | --- |
| Intercept | 0.580 (0.121)*** | 0.575 (0.121)*** |
| NACC | 0.140 (0.063)* | 0.135 (0.064)* |
| IPFC | -0.147 (0.057)* | -0.123 (0.058)* |
| Age | 0.105 (0.086) | 0.104 (0.086) |
| Gender | -0.084 (0.172) | -0.077 (0.172) |
| Variance Component | A – Estimate | B – Estimate |
| $\text{Var}(\pi_{0ij})$ | 0.267 | 0.267 |
| $\text{Var}(\pi_{1ij})$ | 0.041 | 0.043 |
| $\text{Var}(\pi_{2ij})$ | 0.024 | 0.024 |
| Fit Statistic | A – Statistic | B – Statistic |
| AIC | 2888.7 | 2888.7 |
| BIC | 2951.5 | 2951.5 |

*Note.* ‘ ’ refers to $p < .10$ , \* refers to $p < .05$ , \*\* refers to $p < .01$ , \*\*\* refers to $p < .001$ . Gender was dummy coded (0 = male, 1 = female). NACC refers to univariate ventral striatum activity; IPFC refers to univariate lateral prefrontal cortex activity. Var() refers to a variance component of a given random effect from the model. Results come from a multilevel logistic regression model, with log-odds of a risky choice as the dependent variable. The ‘A’ column references results using an IPFC mask thresholded at .25; the ‘B’ column references results using an alternate, more conservative mask (thresholded at .50) that covered less cortical area.

**Figure 5.**
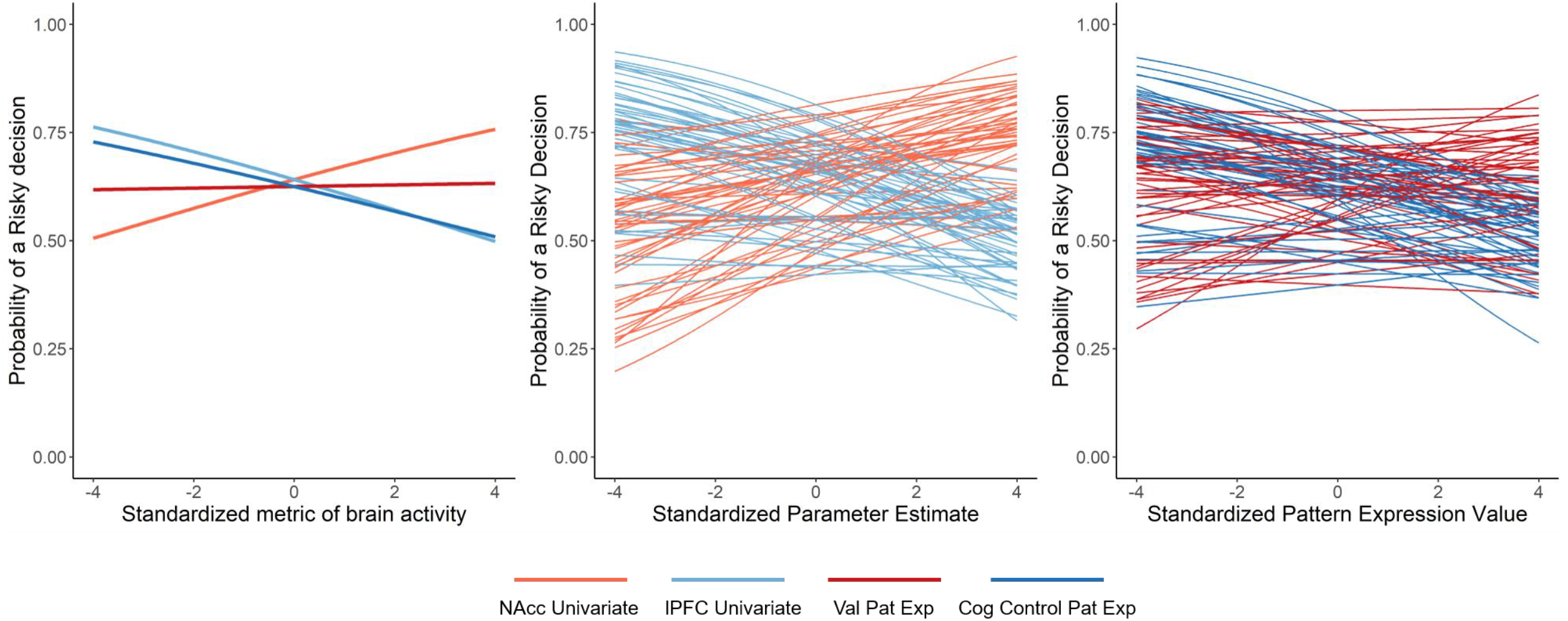
Associations between univariate and multivariate brain activity and probability of making a risky decision *Note.* ‘NAcc Univariate’ refers to the trial-level univariate NAcc activity estimates; ‘lPFC Univariate’ refers to the trial-level univariate lPFC activity estimates; ‘Cog Control Pat Exp’ refers to trial-level cognitive control pattern expression estimates; ‘Value Pat Exp’ refers to trial-level value-based pattern expression estimates. Fixed effects of brain activity metrics from both models are shown in the left panel. Subject specific random effects of associations between risky decision-making probability and univariate NAcc and univariate lPFC activity are depicted in the middle panel. Subject specific random effects of associations between risky decision-making probability and, value-based pattern expression, and cognitive control pattern expression values are depicted in the right panel.

### Classic

Using the classic system-based model, we found that trial-level univariate NAcc and lPFC activity were independently associated with decision tendencies in the yellow light game in a manner consistent with theory. Within-person increases in NAcc activity were associated with an increased likelihood of making a risky decision, whereas within-person increases in lPFC activity were associated with a decreased likelihood of making a risky decision. The magnitudes of the effects were comparable: a one unit increase in NAcc activity was associated with a 15.03% increase in the expected odds of making a risky decision, while a one unit increase in lPFC activity corresponded with a 13.67% decrease in the expected odds of making a risky decision (calculated using coefficients reported in Table 2, Column A). Notably, these results remained highly similar when using an alternate, more conservative lPFC mask (results still significant, same direction, comparable effect sizes; Table 2, Column B). Age did not interact with either NAcc or lPFC activity (Table 4, Column A).

### Switchboard

Results from the switchboard system-based model are summarized in Table 3. These results are partially consistent with system-based theories, in that cognitive control pattern expression estimates were significantly associated with risky decision-making on the YLG. A one unit increase in cognitive control pattern expression corresponded with a 11.57% decrease in the expected odds of making a risky decision (obtained from Table 3, Column A). Sensitivity analyses indicated this effect was robust to variations in neural signatures (e.g., when using uniformity and association maps, unique voxels in uniformity maps, see Table 3, Columns B-C) and these effects were not present when using theoretically orthogonal neural signatures (‘vision’ and ‘auditory’, see Table 5). Collectively these results indicate that multivariate pattern-based activity related to cognitive control encodes meaningful information about risk-taking tendencies. Simultaneously, and inconsistently with system-based theories, value-based pattern expression estimates were not significantly associated with risky decision propensities (coefficient: .051, (SE = .048, ns), from Table 3B). To ensure our value-based results were not driven via the selection of an erroneous pattern, we re-ran analyses using Neurosynth’s ‘reward’ term and obtained near-identical results (reward coefficient: .043 (SE = .048, ns)). Importantly, multivariate patterns and univariate activity metrics were modestly correlated (correlations ranged between approximately .09 and .4), and overlap between the lPFC and cognitive control multivariate signature—the only system that was significant in both model variants—was minimal (only 7.9% of the voxels in the cognitive control signature were also present in the lPFC mask). We reiterate here that univariate activity metrics and multivariate pattern expression scores represent different aspects of brain activity, and these descriptive statistics emphasize this point. As with the classic model, age did not interact with either value or cognitive control patterns.

**Table 3.** Log-odds of risky choice from within-subjects analysis of switchboard system-based models

| Term | A – Estimate | B – Estimate | C – Estimate |
| --- | --- | --- | --- |
| Intercept | 0.581 (0.123)*** | 0.510 (0.122)*** | 0.619 (0.122)*** |
| Value PE | 0.051 (0.048) | 0.008 (0.054) | -0.004 (0.052) |
| Cognitive Control PE | -0.123 (0.047)** | -0.119 (0.060)* | -0.154 (0.055)** |
| Age | 0.113 (0.087) | 0.134 (0.079)' | 0.122 (0.087) |
| Gender | -0.104 (0.181) | 0.054 (0.181) | -0.176 (0.174) |
| Variance Component | A – Estimate | B – Estimate | C – Estimate |
| Var($\pi_{0ij}$) | 0.259 | 0.253 | 0.284 |
| Var($\pi_{1ij}$) | 0.000 | 0.006 | 0.010 |
| Var($\pi_{2ij}$) | 0.003 | 0.039 | 0.022 |
| Fit Statistic | A – Statistic | B – Statistic | C – Statistic |
| AIC | 2894.3 | 2892.6 | 2888.1 |
| BIC | 2957.1 | 2955.4 | 2950.9 |

*Note.* ' refers to $p < .10$ , \* refers to $p < .05$ , \*\* refers to $p < .01$ , \*\*\* refers to $p < .001$ . Value PE refers to value-based pattern expression; Cognitive Control PE refers to cognitive control pattern expression. Var() refers to a variance component of a given random effect from the model. Results come from a multilevel logistic regression model, with log-odds of a risky choice as the dependent variable. The 'A' column references results using Neurosynth association maps for pattern expression analysis; the 'B' column references results using Neurosynth uniformity maps; the 'C' column references results from a pattern expression analysis using only unique voxels among the value and cognitive control Neurosynth uniformity maps (i.e., common voxels between the two masks were removed).

**Table 4.** Models testing interactions with age

| Term | A – Estimate | B – Estimate |
| --- | --- | --- |
| Intercept | 0.583 (0.121)*** | 0.508 (0.122)*** |
| NACC (A) \| Value PE<br>(B) | 0.120 (0.063)' | 0.007 (0.054) |
| IPFC (A) \| Cognitive<br>Control PE (B) | -0.139 (0.057)* | -0.117 (0.061)' |
| Age | 0.107 (0.087) | 0.120 (0.086) |
| Gender | -0.088 (0.171) | -0.056 (0.182) |
| NACC (A) \| Value PE<br>(B) x Age | -0.090 (0.063) | 0.004 (0.056) |
| IPFC (A) \| Cognitive<br>Control PE (B) x Age | 0.045 (0.057) | 0.019 (0.062) |
| Variance Component | A – Estimate | B – Estimate |
| $\text{Var}(\pi_{0ij})$ | 0.266 | 0.252 |
| $\text{Var}(\pi_{1ij})$ | 0.033 | 0.006 |
| $\text{Var}(\pi_{2ij})$ | 0.022 | 0.036 |
| Fit Statistic | A – Statistic | B – Statistic |
| AIC | 2890.8 | 2896.5 |
| BIC | 2965.0 | 2970.7 |

*Note.* ' refers to $p < .10$ , \* refers to $p < .05$ , \*\* refers to $p < .01$ , \*\*\* refers to $p < .001$ . Gender was dummy coded (0 = male, 1 = female). NACC refers to univariate ventral striatum activity; IPFC refers to univariate lateral prefrontal cortex activity. Var() refers to a variance component of a given random effect from the model. Results come from a multilevel logistic regression model, with log-odds of a risky choice as the dependent variable. The 'A' column references results from the classic model (IPFC threshold = 0.25); the 'B' column references results from the switchboard model (association maps). In order to be concise, differing terms for each model (any term involving a metric of brain activity) are included in the same line of the first column, separated by '|'.

**Table 5.** Log-odds of risky choice from models with additional Neurosynth patterns to gauge uniqueness of cognitive control pattern expression findings

| Term | Estimate |
| --- | --- |
| Intercept | 0.569 (0.120)*** |
| Vision PE | 0.096 (0.054)' |
| Auditory PE | 0.075 (0.050) |
| Age | 0.072 (0.083) |
| Gender | -0.072 (0.168) |
| Variance Component | Estimate |
| $\text{Var}(\pi_{0ij})$ | 0.292 |
| $\text{Var}(\pi_{1ij})$ | 0.027 |
| $\text{Var}(\pi_{2ij})$ | 0.011 |
| Fit Statistic | Statistic |
| AIC | 2888.4 |
| BIC | 2951.2 |

*Note.* ' refers to $p < .10$ , \* refers to $p < .05$ , \*\* refers to $p < .01$ , \*\*\* refers to $p < .001$ . Gender was dummy coded (0 = male, 1 = female). The alternate maps correspond to the terms listed in the table ('Vision', 'Auditory'). 'PE' refers to pattern expression. Association maps for each term were used. Var() refers to a variance component of a given random effect from the model. Results come from a multilevel logistic regression model, with logodds of a risky choice as the dependent variable.

We conducted *post-hoc* analyses to interrogate the lack of a relationship between value-based pattern expression estimates and risky behavior. We first examined whether multivariate signatures within the NAcc were associated with behavior given that univariate signals within this region were, operating under the logic that value-based patterns may be more localized to a given region than cognitive control. We re-ran the pattern expression analyses with the value-based neural signature, but this time only included voxels in the NAcc in our mask. Again, this analysis showed a non-association between value-based pattern expression scores in the NAcc and risky decision-making (coefficient = 0.000, *ns*). Given this result and the nature of pattern expression analysis, it was puzzling why univariate activity in the NAcc tracked with behavior (especially when considering a bulk of the pattern was comprised of NAcc voxels, see Figure 3), but value-based signatures—even if localized to the NAcc—did not. This led us to believe that perhaps it was the homogeneity of multivariate activity in NAcc that related to risky decision tendencies. Multivariate patterns necessarily encode spatial variability, but it could that homogeneity or uniformity of activity are more strongly predictive of behavior, suggesting that pattern expression estimates that inherently capture this variability may be poor predictors of behavior. To test this, we re-extracted multivariate patterns from the NAcc and lPFC and computed Gini coefficients for each region for each trial. Traditionally used in macroeconomics but recently applied in neuroscience (Guassi Moreira, McLaughlin, & Silvers, 2019; Guest & Love, 2017), Gini coefficients in this context can describe the extent to which brain activity in a given region is homogenous (uniform) or heterogeneous. Indeed, as shown in Table 6, a lower Gini coefficient in the NAcc (i.e., more uniform activation) was associated with an increased propensity to take risks on the YLG, suggesting a strong, one-dimensional encoding of value signatures during decision-making.

**Table 6.** Models with predicting decision activity from trial-level Gini coefficients

| Term | Estimate |
| --- | --- |
| Intercept | 0.629 (0.121)*** |
| NACC Gini | -0.115 (0.048)* |
| IPFC Gini | 0.057 (0.047) |
| Age | 0.080 (0.088) |
| Gender | -0.198 (0.172) |
| Variance Component | Estimate |
| $\text{Var}(\pi_{0ij})$ | 0.270 |
| $\text{Var}(\pi_{1ij})$ | 0.009 |
| $\text{Var}(\pi_{2ij})$ | 0.004 |
| Fit Statistic | Statistic |
| AIC | 2894.1 |
| BIC | 2956.9 |

*Note.* ‘ refers to $p < .10$ , \* refers to $p < .05$ , \*\* refers to $p < .01$ , \*\*\* refers to $p < .001$ . Gender was dummy coded (0 = male, 1 = female). Var() refers to a variance component of a given random effect from the model. Results come from a multilevel logistic regression model, with log-odds of a risky choice as the dependent variable.

### Between Subject Results

Results from between subject analyses indicate that none of the between-subject brain activity metrics (univariate or pattern based) were related to proportion of risky decisions (univariate NAcc: *b* = 0.223, SE = 0.316, *p* > .250; univariate lPFC: *b* = −0.209, SE = 0.270, *p* > .250; value pattern expression: 8.725e-5, SE = 1.823e-4, *p* > .250; cognitive control pattern expression: −1.435e-4, SE = 1.219e-4, *p* = .245). A traditional brain mapping (mass univariate) analysis of the YLG showed significant anterior cingulate cortex (ACC) activity for the ‘Go > Stop’ contrast in addition to significant amygdala and dorsal striatal activity (Figure 6; Table 7).

**Table 7.** Brain regions which showed significant activation Go > Stop and Stop > Go.

| Region |  | x | y | z | Z | k |
| --- | --- | --- | --- | --- | --- | --- |
| *Go > Stop* |  |  |  |  |  |  |
| Occipital Pole | L | -18 | -90 | 0 | 7.51 | 13177<sup>a</sup> |
| Striatum | R | 10 | 6 | 8 | 4.22 | <sup>a</sup> |
| pSTS | L | -52 | -36 | 28 | 5.18 | 491 |
| ACC | R | 12 | 10 | 42 | 5.15 | 406 |
| Amygdala | L | -30 | -10 | -14 | 5.60 | 372 |
| Amygdala | R | 30 | -12 | -12 | 5.86 | 229 |
| Insular Cortex | L | -56 | 4 | 12 | 4.49 | 216 |
| SPL | R | 24 | -52 | 56 | 5.34 | 146 |
| *Stop > Go* |  |  |  |  |  |  |
| TPJ | R | 50 | -56 | 42 | 4.97 | 566 |
| TPJ | L | -46 | -60 | 46 | 4.52 | 204 |

*Note.* R refers to right and L refers to left. x, y, and z refer to MNI coordinates; Z refers to the z-statistic at those coordinates (local maxima); pSTS refers to posterior superior temporal sulcus; ACC refers to anterior cingulate cortex; IOG refers to inferior occipital gyrus; ACC refers to anterior cingulate gyrus; SPL refers to superior parietal lobule; TPJ refers to temporoparietal junction. Regions that share the same superscript are part of the same cluster.

**Figure 6.**
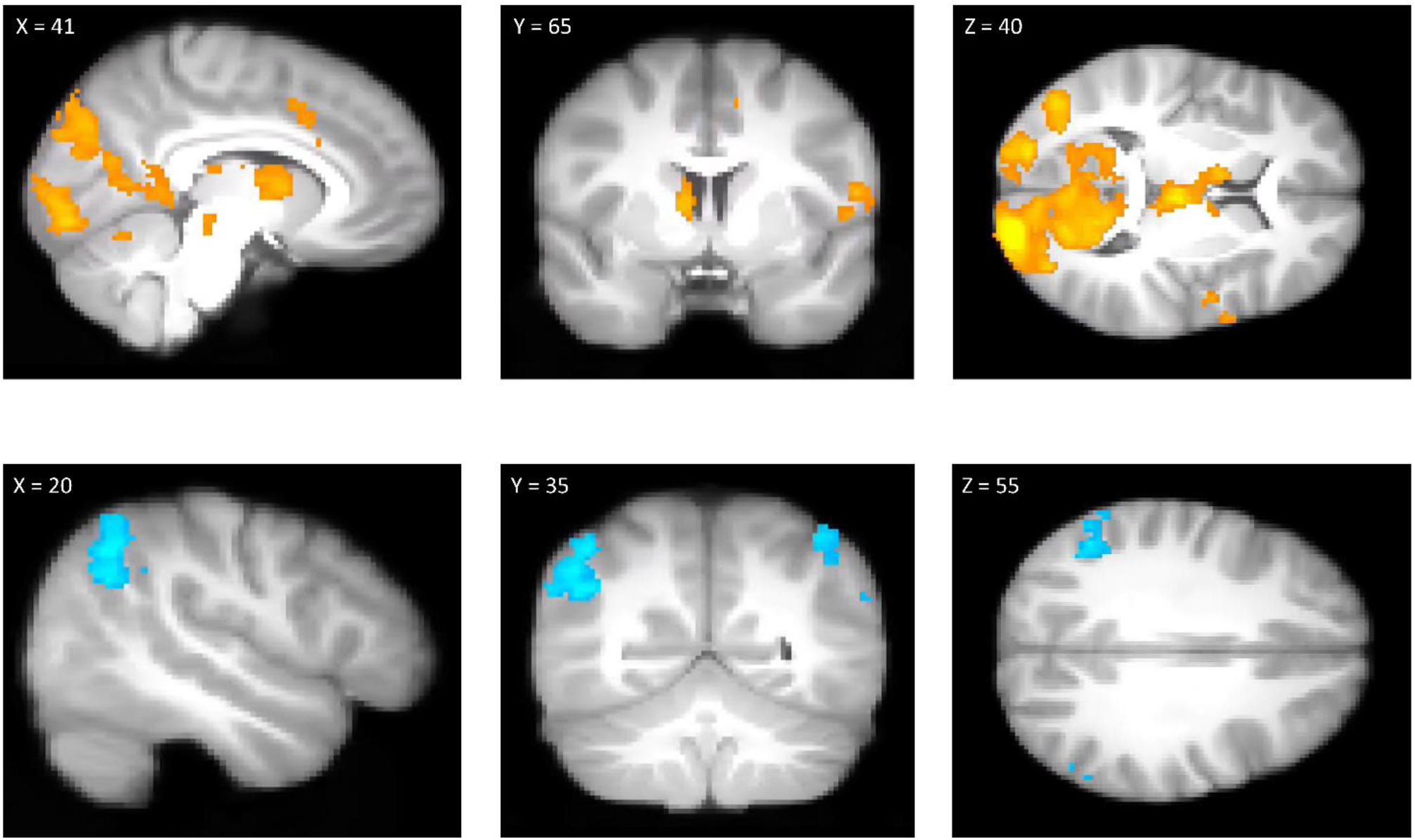
Results from the Go > Stop (top row) and Stop > Go (bottom row) contrasts. *Note.* XYZ refer to voxel coordinates in MNI standard space. Clusters rendered here were obtained using a cluster-defining-threshold of Z = 3.1, correct for multiple comparisons at *p* < .05 using Random Field Theory. Clusters are rendered on ‘bg_image’, FSL’s average of all subjects’ high resolution anatomical images.

## Discussion

The current study employed a brain modeling philosophy (Kragel et al., 2018) in conjunction with within-subject multilevel logistic regression to test system-based theories of adolescent decision-making. In doing so, we also expanded upon traditional neuroscientific implementations of system-based theories by examining the role of multivariate neural signals. We found that features of brain activity predicted behavior in a manner consistent with system-based theories. We observed this in two variants of the model—a classic implementation assuming modularity among ROIs, and a novel variant that included information for multivariate signatures. These findings have a number of ramifications for neuroscientific models of adolescent decision-making.

### Implications for System-Based Theories of Adolescent Decision-Making

We observed that value-based and cognitive control systems generally predicted behavior in a manner consistent with system-based theories: univariate estimates of NAcc and lPFC activity were directly and inversely, respectively, associated with the probability of making risky decision, while cognitive control pattern expression was predictive of a reduced likelihood to make a risky decision. Our between subject analyses, along with traditional mass univariate brain mapping analyses, failed to show any such trends. Two broad conclusions follow from these results. First, these findings support the utility of brain modeling techniques for testing system-based theories of decision-making in developmental neuroscience and beyond, reinforcing that brain modeling and brain mapping philosophies are not simply inverse functions of the other that yield equivalent results (Kragel et al., 2018). Second, and perhaps more importantly, these results suggest features of system-based theories of decision-making carry evidentiary value, despite compelling arguments to the contrary (Pfeifer & Allen, 2012). The present findings suggest that system-based theories may offer interim frameworks for relatively young fields such as cognitive neuroscience as they continue to incrementally extend theory on the basis of novel evidence (Baddeley, 2012; Pfeifer & Allen, 2016). Even as these theories are eventually replaced by stronger accounts that consider more nuanced relationships between cognitive control and value systems as well as other biological influences (Davidow et al., 2018; Harden et al., 2017), their use as ‘baseline’ models may actually facilitate novel theoretical insights so long as they are not subscribed to too rigidly. Furthermore, that features of system-based models have evidentiary value is not tantamount to saying they are optimal (indeed, a comparison of model fit statistics between system-based models and empty or covariate-only models in the present study suggests otherwise), but rather points to the need to develop and test more nuanced quantitative models of the neurodevelopment of decision-making behavior. Relatedly, we failed to observe interactions between brain activity metrics and age, a major tenet of developmental system-based models. This null finding underscores our call for greater nuance in quantitative models of decision-making neurodevelopment: clear age-related behavioral differences in risk-taking behavior (Defoe, Dubas, & Romer, 2019; Duell et al., 2017) are necessarily encoded in brain activity, yet current modeling approaches have been unable to consistently link age differences in the association between neurobiology and behavior.

### Modularity and Population Coding in System-Based Theories of Decision-Making

On a more granular level, our results speak to two long-discussed concepts in neuroscience: modularity and population coding (Erickson, 2001). These two concepts are respectively reflected by univariate and multivariate analyses in neuroimaging data. Most system-based theories of adolescent decision-making originate from disciplines within psychological science that espoused modularity at the *psychological level* (Steinberg et al., 2008). Although it is not a given that psychological modularity necessitates neural modularity, this assumption has been preserved in many neuroscientific implementations of system-based theories (Shulman et al., 2016; Strang et al., 2013), despite evidence in adults that multivariate patterns reflect meaningful information about decision-making (Hampton & O’Doherty, 2007). Our univariate and multivariate results, somewhat surprisingly, respectively provide support for *both* modularity and population coding (Cosme & Lopez, 2020) – specifically, results revealed that univariate NAcc and lPFC activity was associated with decision behavior, and also that pattern expression of a multivariate cognitive control (but not value) signature predicted decisions. The former (evidence of modularity) is surprising, given the limited support for neural modularity that prior studies have found (Erickson, 2001), whereas the latter (population coding) is notable because no prior studies, to our knowledge, have found evidence of such in the context of brain modeling decision behavior (i.e., using multivariate metrics to model behavior/cognition).

These findings carry notable implications. Although our results suggest that modularity may be a feature of adolescent decision behavior, we cannot conclude with certainty what activation in those modules (i.e., NAcc and lPFC) reflects. While such activation could index computations related to value and cognitive control, respectively, it is more difficult to infer psychological processes from ROI-based activity than from multivariate signatures, which tend to be more specific in what they reflect (Poldrack, 2006; Wager et al., 2013). Our data are roughly consistent with an amplifier model, which would allow for reconciliation of our classic and switchboard results. In such a model, multivariate patterns may code for the psychological process of interest and the modules observed here act as ‘volume’ knobs that amplify their magnitude. In other words, the multivariate patterns code for a given psychological process whereas the univariate activity of the modules controls the intensity of the process. Indeed, such multidimensional coding schemes appear to support decision behaviors in monkeys (Zhang, Chen, & Monosov, 2019), and similar findings from human samples in other domains (eating behavior) further hint at the neural plausibility of a modular-population hybrid scheme (Cosme & Lopez, 2020). Further work could also examine whether there is a qualitative shift between coding schemes across development (Gee et al., 2013). Though we found no such evidence in our own data, future work could broaden age ranges to include young children and adults to determine the extent to which system-based models explain decision making at different developmental stages. Overall, it is clear that additional work is needed to characterize the relative contributions of neural modules and population codes in system-based theories of decision-making, involving the use of different behavioral tasks, different multivariate signatures, and evaluation of decision behaviors in different contexts.

### Building on System-Based Theories of Adolescent Decision-Making

Our findings suggest system-based models provide at least some explanatory utility, but it is critical that future work improves upon existing models in several key ways. One future step will be to determine the algebraic form of influential system-based theories. As we noted before, existing neuroscientific system-based theories of human decision-making in linguistic terms without specifying a computational model (i.e., they are explained qualitatively, instead of with an algebraic equation). This means one could posit a number of algebraic forms that satisfy qualitative requirements of system-based theories that each carry very different implications. We assumed a linear relationship between the log odds of a risky choice and metrics of brain activity, but an alternate statistical model may be more appropriate. Future studies could test a set of candidate algebraic formulations of system-based decision-making theories (e.g., estimating latent value and linking to decision likelihoods, etc.). This could facilitate cross-study and cross-discipline comparison by setting an objective framework that supports falsifiability. Future work must also directly address our null findings involving value-based pattern expression values. While we tested ‘reward’ and ‘value’ patterns and obtained null results with both, it is possible an alternative untested pattern computed in a different manner (i.e., not relying on meta-analytic maps) would yield positive results. To rigorously test this possibility, we recommend future studies systematically create and test patterns that vary iteratively on psychological processes relevant for system-based theories (Chang et al., 2015; Wager et al., 2013). This process should also involve understanding how such maps change with development, as it may be unrealistic to assume a reward signature derived in one age group is readily applicable to all ages. Taking such an approach would also have the benefit of providing insights into what specific psychological features these patterns encompass – for example, by examining subcomponents of cognitive control (e.g., working memory). Finally, it is worth noting that interactions between each system and age were null, defying a core feature of system-based theories, suggesting that more bottom-up exploratory work is needed to better understand how the dynamic potency of each system changes with age. Ideally, such work would involve repeated sampling at both the decision- and subject-level (i.e., longitudinal assessments).

### Limitations and future directions

The current study has several limitations that point directly to future directions in this line of research. The first is that the effect sizes found from key results are somewhat modest. Though not a traditional ‘limitation’ per se, this points to the possibility that other untested computational signals in the brain may also contribute to decision behavior. Another limitation is that the present results were obtained in a single, moderately-sized sample and ought to be replicated (Helmer et al., 2020; Marek et al., 2020). That said, our concerns about samples size are partially assuaged by the fact that we leveraged multilevel models to maximize statistical power when examining brain-behavior associations (Schoeneberger, 2016). Three additional limitations also exist regarding generalizability. First, in terms of adolescent decision-making, prior work shows adolescent decision behaviors are prone to tremendous diversity across the world and even within individuals (Steinberg et al., 2017), forcing us to consider that these results, even ignoring other limitations, may not reflect a ‘common ground truth’ among all humans or even within a single human (to the extent such a ‘ground truth’ actually exists). Second, it is possible that a different pattern of results would emerge for decision behaviors in other contexts (e.g., moral, financial decisions). Third, it is unclear whether these findings are specific to adolescence or generalize to general decision-making processes across the lifespan. A final limitation is that we did not compare our implementation of system-based theories to alternative theories. While this is mainly because system-based theories have dominated the field and alternative approaches have been relatively atheoretical (Pfeifer & Allen, 2012, 2016), we look forward to future work aimed at rigorously comparing alternate explanations.

### Concluding Remarks

System-based theories of adolescent decision-making have drawn tremendous scholarly interest, yet the veracity of their neuroscientific implementation has been the subject of much debate. This investigation was the first to our knowledge to test system-based theories of adolescent decision-making using a methodological approach that is more consistent with the core tenets of such theories (i.e., brain modeling). We found evidence that system-based theories are indeed predictive of adolescent risk-taking behaviors, showing that univariate and multivariate brain activity metrics of cognitive control and value-based processes predict trial-by-trial risky decision tendencies. We did not, however, observe evidence that these neural systems interacted with age, at odds with a key element of system-based theories. Overall, this work contributes knowledge about the neural bases of adolescent decision behavior.

## Acknowledgments

Preparation of this manuscript was supported by a National Science Foundation Graduate Research Fellowship (2016220797) and a National Institutes of Health Predoctoral T32 Fellowship to JFGM, and generous funds from the UCLA academic senate, UCLA Hellman Fellows Fund, and an NSF CAREER grant (1848004), to JAS. We thank Drs. Jennifer Pfeifer and Shannon Peake for sharing the Yellow Light Game with us. We appreciate the efforts of Austin Blake, Alejandra Delgado, Zoey Dew, Milagro Escobar, Elizabeth Gaines, Agatha Handojo, Ciara Mandich, Nora Ngo, Sakina Qadir, Emily Towner, Claire Waller, and Tarran Walter in participant recruitment and data collection. We are also grateful for Maria Calderon Leon’s contributions to a relevant literature search. We are grateful for reflections on the study concept from Dr. Jamie Feusner, Dr. Andrew Fuligni, Dr. Adriana Galván, Amanda E. Baker, Zoe Guttman, and Dr. Monica Rosenberg’s Cognition, Attention, and Brain Lab at the University of Chicago.

## Extended Data

**Table 1-1.** Head motion statistics

|  | *n* Volumes > 0.9mm<br>(mean, SD) | Most extreme displacement<br>(mean, SD) | % *n* Volumes < 0.9mm | % Extreme displacements<br>< 1.8mm |
| --- | --- | --- | --- | --- |
| Run 1 | 0.0816, 0.5714 | 0.3270, 0.2894 | 97.96% | 100% |
| Run 2 | 0.2653, 0.8845 | 0.5013, 0.7125 | 87.76% | 95.92% |
| Run 3 | 0.3265, 1.1617 | 0.4342, 0.4876 | 91.84% | 95.92% |

*Note.* ‘*n* Volumes > 0.9mm’ reflects the average number of volumes exceeding the 0.9mm frame displacement threshold across subjects; ‘Most extreme displacement’ reflects the average of the most extreme frame displacements across subject. ‘% *n* Volumes < 0.9mm’ indicates the percentage of subjects whose data did not have a single volume exceeding our 0.9mm frame displacement cut-off. ‘% Extreme displacements < 1.8mm’ indicates the percentage of subjects whose most extreme frame displacement value did not exceed 1.8mm, corresponding to twice the value of our 0.9 mm threshold.

**Figure 1-1.**
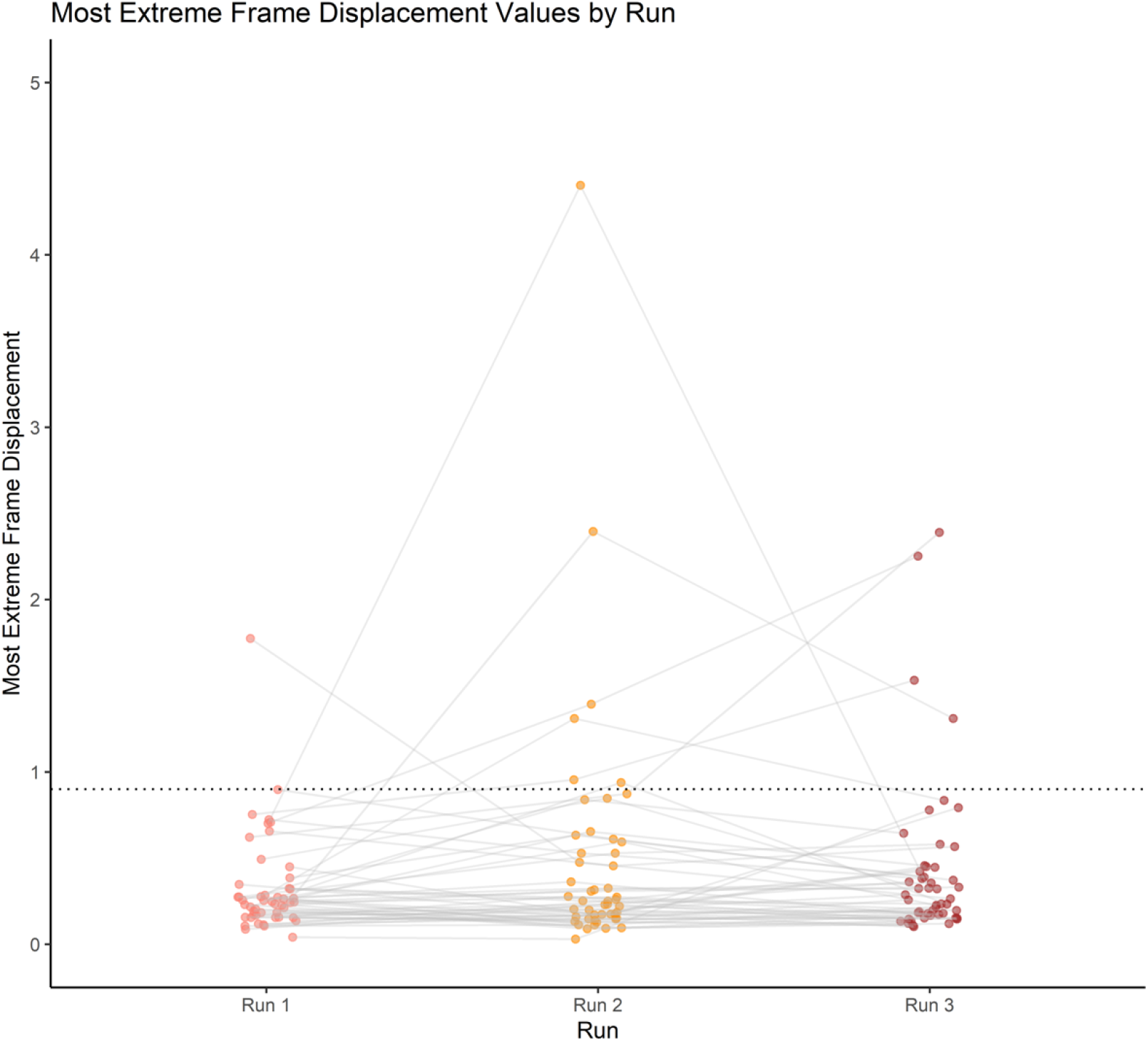
Most extreme frame displacements *Note.* Values are randomly jittered along the x-axis. Frame displacement values are in millimeter units. The dotted line corresponds to our frame displacement cutoff of 0.9.

